# Novel Proteomic changes in Yeast Mitochondria provide insights into mitochondrial functioning upon over-expression of human p53

**DOI:** 10.1101/746412

**Authors:** Archana Kumari Redhu, Jayadeva Paike Bhat

**Affiliations:** Laboratory of Molecular Genetics, Department of Biosciences and Bioengineering, Indian Institute of Technology-Bombay, Powai. Mumbai-400076

**Author notes:** Corresponding author: Jayadeva Paike Bhat.

**Keywords:** p53, *Saccharomyces cerevisiae*, Mitochondria, Apoptosis, Proteomics

## Abstract

Cancer cells display enhanced glycolytic activity and impaired oxidative phosphorylation even in the presence of adequate oxygen (Warburg effect). Mitochondrial physiology is a promising hit target for anti-cancer therapy because of its key role in Warburg effect and activating apoptosis in mammalian as well as yeast cells. Over-expression of human p53 in *S.cerevisiae* leads to cell cycle arrest and apotosis. In the present work we show that how *S.cerevisiae* escapes from p53 induced apoptosis in fermentable carbon source, whereas in case of non-fermentable carbon source this phenomenon is not observed. To shed the light on this aspect we performed a quantitative proteomic analysis of yeast mitochondria isolated from the cells grown on sucrose (fermentation) and glycerol (respiration) with and without p53 over-expression. Through this approach, we identified a total dataset of 1120 proteins with 1% FDR, of which 239(133+106) proteins are differentially experssed in both conditions. Interestingly, we observed that after over-expression of p53 in sucrose grown yeast cells, a complete set of pentose phosphate pathway (PPP) enzymes is up-regulated in the mitochondria that leads to enhanced mitochondrial NADPH production and ROS quenching. Increased association of a hexose transporter (HXT6) and a hexokinase (HXK2) with the mitochondria of fermenting yeast cells upon over-expression of p53, may direct glucose towards PPP inside the mitochondria. In conclusion, our results provide the evidence that up-regulated PPP inside the mitochondria is a key to evade apoptosis by *S.cerevisiae* upon p53 over-expression.

## Introduction

Warburg effect and Crabtree effect are the two most well studied and important metabolic regulatory features of yeast *S.cerevisiae* that come into play during its rapid growth on sufficient glucose as a sole carbon source (1,2,3). These two regulatory features are recapitulated in human cancer cells and were first identified in the 1920s (4, 5). Much later, it was observed that these features are also exhibited by rapidly proliferating human cells (6), implying that the regulatory phenomenon underlying rapid cell proliferation using glucose as the carbon source is evolutionarily conserved from microorganisms to humans. Interestingly, that regulatory mechanisms of microbial metabolism can be used as a stepping stone for the understanding of tumour cell metabolism was proposed as early as the 1930s (7). In the past decade, these two regulatory phenomena have been intensely studied both in bacteria and yeast under the term overflow metabolism (8,9). There is now compelling evidence to suggest that genes that are preferentially expressed in the tumour are conserved in microbial species (10) and may be responsible for the overflow metabolism. Thus, it is not surprising that a variety of experimental approaches have been used in yeast to unravel the molecular mechanisms of aerobic glycolysis as well as Crabtree effect (11,12).

Apoptosis is yet another regulatory phenomenon that has been conserved from yeast to humans (13). Escape from apoptosis is one of the hallmarks of tumour cells (14). It has been demonstrated that the expression of p53 and Bax induces apoptosis in yeast (15,16). Mitochondria play a major role, both in yeast and mammalian apoptosis (17). Moreover, yeast cells lacking HXK2 constitutively accumulate Ras in mitochondria and are hypersensitive to AIF1 dependent apoptosis in response to inducing agents such as H_2_O_2_ and acetic acid (18). In transformed cells, 70% of HXK2 is bound to mitochondria via its interaction with mitochondria membrane protein VDAC (19, 20). Over-expression of p53 in a crab tree positive yeast i.e. *S.cerevisiae*, causes apoptosis only when cells are grown in glycerol, a non-fermentative medium but not in a fermentative medium. In contrast, in *K. lactis*, Crabtree-negative yeast, over-expression of p53 causes apoptosis in both fermentative as well as non-fermentative yeast (21). This study for the first time demonstrated that p53 induced apoptosis in yeast species is in some way connected through Crabtree effect.

The aim was to investigate how *S. cerevisiae* escapes from p53 induced apoptosis only when cells are grown in a fermentable carbon source. The approach used was to achieve a more detailed characterization of the yeast mitochondrial proteome to study differential expression of proteins in fermentable or non-fermentable conditions with or without p53 over-expression. After analysis and identification of the detected mitochondrial proteins, we observed that the expressions of pentose phosphate pathway (PPP) enzymes, which catalyse the irreversible steps of PPP, were higher only in mitochondria isolated from sucrose grown cells upon over-expression of p53. On the other hand, mitochondria isolated from glycerol-grown cells upon over-expression of p53 did not follow such a pattern. Thus, it appeared that the cells grown in sucrose even upon over-expression of p53 do not undergo apoptosis most probably because of higher NADPH content provided by PPP pathway operated inside their mitochondria. Here, in continuation with our efforts we investigated a novel role of HXK2 and HXT6 in p53 induced escape from apoptosis in the cells grown in sucrose, more interestingly we observed that apoptosis induced by lack of Hxk2 might be because of reduced amount of G6PD inside the mitochondria. The above results clearly indicate that the entry of PPP enzymes into mitochondria can prevent p53 induced apoptosis.

## Material and Methods

### Materials

Geneticin (G418) and NADP monosodium salt were purchased from Calbiochem®. G6PDH enzyme, glutathione reductase (GR), Glucose-6-phosphate, 6phosphogluconate (6PG), EDTA, Sorbitol, L-glutathione oxidised disodium salt, Phenylmethanesulfonyl fluoride (PMSF), β-mercaptoethanol, protease inhibitor cocktail was purchased from Sigma. Zymolase-20T was purchased from MP Biomedicals, LLC (India).

Antibodies against the following proteins/epitopes were purchased from the mentioned sources: p53 (Santa Cruz-Biotechnology (FL-393, F#2513); G6PDH (A-9521 Sigma); HA Probe (Y-11, E#1712 Sigma); alkaline phosphatase conjugated IgG (A-3687 Sigma); VDAC-1(16G9E6BC4 Invitrogen), β-actin (mABGEa, NB100-74340SS Novus Bio). The oligonucleotides used in the study, as listed in Supplementary Table-I, were procured from Sigma Life Sciences, India.

### Strain and Culture conditions

Yeast strains used in this study are listed in Supplementary Table-II. All the strains were grown in minimal medium containing 0.67% (w/v) yeast nitrogen Base (Difco) and ammonium sulphate mixture (1:3), 0.05% (w/v) of the amino acid mixture (complete or drop out). The carbon sources were 3% (v/v) glycerol plus 2% (v/v) potassium lactate or 0.2% (w/v) sucrose or 2% (w/v) galactose or 2% (w/v). All gene disruptions were performed with *KanMX4* cassette amplified from the pUG6 vector using appropriate primers. Yeast strain deletion transformants were selected on YEPD plates containing G418 at a final concentration of 200μg/ml. Plasmid pYM14 was used for C-terminal 6xHA tagging of 6PGDH. Plasmid pYM28 was used for C-terminal EGFP tagging of HXK-2.

### Spotting Assay

The cells were grown in minimal medium containing either Glycerol or Sucrose as main carbon source. The cells were harvested at mid-log phase and O.D._600_ of 1.0 was achieved for all the cultures by diluting the samples with sterile distilled water. 4 μl of fivefold serial dilution was spotted on YEPD and plates containing minimal media. Growth differences were recorded following incubation of plates for 48-72hrs at 30°C.

### Isolation of Mitochondria from *S. cerevisiae*

wtp53 strain was grown in sucrose and glycerol lactate medium separately. For p53 over-expression mid-exponential phase grown cells were induced with 2% galactose and were incubated further for 6-8 hours. Another set of cells induced with an equal amount of water were used as control. Cells were then harvested by centrifugation for 5 minutes at 3000g. Pellet was rinsed with 10 mM EDTA, and resuspended in 1.2 M sorbitol buffer (1.2 M sorbitol, 50 mMTris/HCl, pH 7.5, 10 mM EDTA, 0.3% β-mercaptoethanol). The cell wall was digested enzymatically with Zymolyase-20T at 1 mg/g of cells at 37°C for 30 min. The spheroplasts were harvested by centrifugation at 1700*g* and were resuspended in ice-cold 0.7 M homogenisation buffer (0.7 M sorbitol, 50 mM Tris/HCl, pH 7.5, 0.2 mM EDTA). The unbroken cells, nuclei, and large debris were pelleted by centrifugation at 2500g. The resulting supernatant was pelleted by centrifugation at 12000*g*. To get 100% pure mitochondria, sucrose density gradient ultracentrifugation was performed where a gradient of sucrose was prepared into a backman ultra-clear centrifugation tube with 1.5ml of 60% sucrose overlayed with 4ml of 32%, 1.5ml of 23% and 1.5ml of 15% sucrose (all w/v) respectively. 3 ml of crude mitochondrial extract was placed on the top of 15% sucrose followed by centrifugation in a Beckman SW41 Ti swinging bucket rotor for 90 minutes on 134000g at 4°C. The intact mitochondrial band at interface of 60/30% was carefully removed and resuspended into SEM buffer (10 mM MOPS/KOH (pH 7.2), 250mM sucrose, 1mM EDTA). Pure mitochondria were pelleted by centrifugation in an MLS-5 rotor for 30 minutes at 10000g (22).

### Digestion Protocols

#### In-Gel digestion for Label-free proteomic analysis by LC-MS/MS

For digestion, 50 μg of mitochondrial protein from all 4 samples were fractionated on SDS PAGE gel followed by staining with Coomassie brilliant blue (CBB) dye; time of separation was kept shorter to reduce the number of gel slices. The gel slices were washed thrice with wash buffer (50% acetonitrile (ACN), 50mM ammonium bicarbonate (ABC) to remove the CBB stain completely. The gel pieces were incubated in a reduction solution (10mM DTT and 100mM ABC buffer) for one hour at 56°C to break the disulphide bonds followed by addition of alkylation solution (50mM iodoacetamide (IAA) in 100 mM ABC buffer and kept in dark for 20 minutes. After washing with ABC buffer, trypsin was added in a ratio of 1:10 (w/w). Gel pieces were kept overnight at 37°C after the addition of trypsin solution (23). Finally, the trypsin-digested samples were further processed using Zip-Tip pipette tips and stored at −20°C and subjected to LC-MS/MS analysis.

#### In-Solution Digestion for Label-free proteomic analysis by LC-MS/MS

To obtain In-Sol trypsin digestion mitochondrial protein samples was transferred to a suspension of 8M urea /100mM NH_4_CO3 followed by two minutes of sonication. To reduce disulphide bonds 5mM TECP and 40mM iodoacetamide (for alkylation) was used for a period of 30 minutes in dark at room temperature. For controlled trypsin digestion ten micrograms of trypsin (Promega) was added to the protein sample (protein to trypsin ratio = 50:1), and digestion was carried out at 37°C overnight (24). C18 clean-up using Sep-Pak columns was performed after digestion of the samples according to the manufactures instructions.

#### LC-MS analysis and Label-free Quantification

LC-MS/MS analysis of purified and digested *S.cerevisiae* mitochondrial lysates was performed on Orbitrap Fusion LC-MS (MININT-CVDFOSE\Fusion) which was connected to an electrospray ion source. The peptides were dissolved in solvent A (0.1% formic acid in 2% acetonitrile and 98% H2O), and then loaded onto a manually packed reverse phase C18 column (PepMap TM RSLC resin C18 75 μm x 50 cm, 2μm particle size, and 100Å) coupled to Thermo EASY-nLC-1200 system. Peptides were eluted from 5% to 95 % solvent B (0.1% formic acid in 90% acetonitrile and 10% H2O) in solvent A at a flow rate of 300 nL min^−1^. For a 180 min gradient, the conditions were as follows: 5-30 % B over 165 min, 30-95% over 5 min, and then held at 95% B for 10 min. Parameters were as follows: for full MS spectra, the scan range was m/z 375-1700 with a resolution of 60,000 at m/z 200. MS/MS acquisition was performed in top speed mode with 3s cycle time. The resolution was 15000 at m/z 200. The intensity threshold was 5000, and the maximum injection time was 30 ms. The AGC target was set to 10000, and the isolation window was 1.2 m/z. Ions with charge states 2+, 3+ and 4+ were sequentially fragmented by higher energy collisional dissociation (HCD) with normalized collision energy (NCE) of 30%. The dynamic exclusion duration was set as 40 s.

#### Analysis of MS Data

Raw files of *S.cerevisiae* mitochondrial protein identification were analyzed by Proteome Discoverer (Thermo Fisher Scientific, version 2.2) and Mascot version 2.2.1 (Matrix Science) software with a percolator (strict FDR value less than 1) against SWISS-PROT (www.expasy.org/sport/), and SGD (www.yeastgenome.org) database. For variable modification methionine oxidation and protein N-terminal acetylation were chosen, and for fixed modification cysteine alkylation by iodoacetamide was chosen. Value of mass errors for parent ion mass was ±10ppm with a fragment ion of ± 0.5Da. Peptide identifications with less than 1% false discovery rate (FDR) were accepted with high confidence (25). Peptides having Mascot score less than 20 were discarded (25). PSMs with high confidence applied at the protein level. Fig.1 represents a schematic workflow for mitochondrial protein isolation, purification, mass spectrometry and analysis.

**Figure 1:**
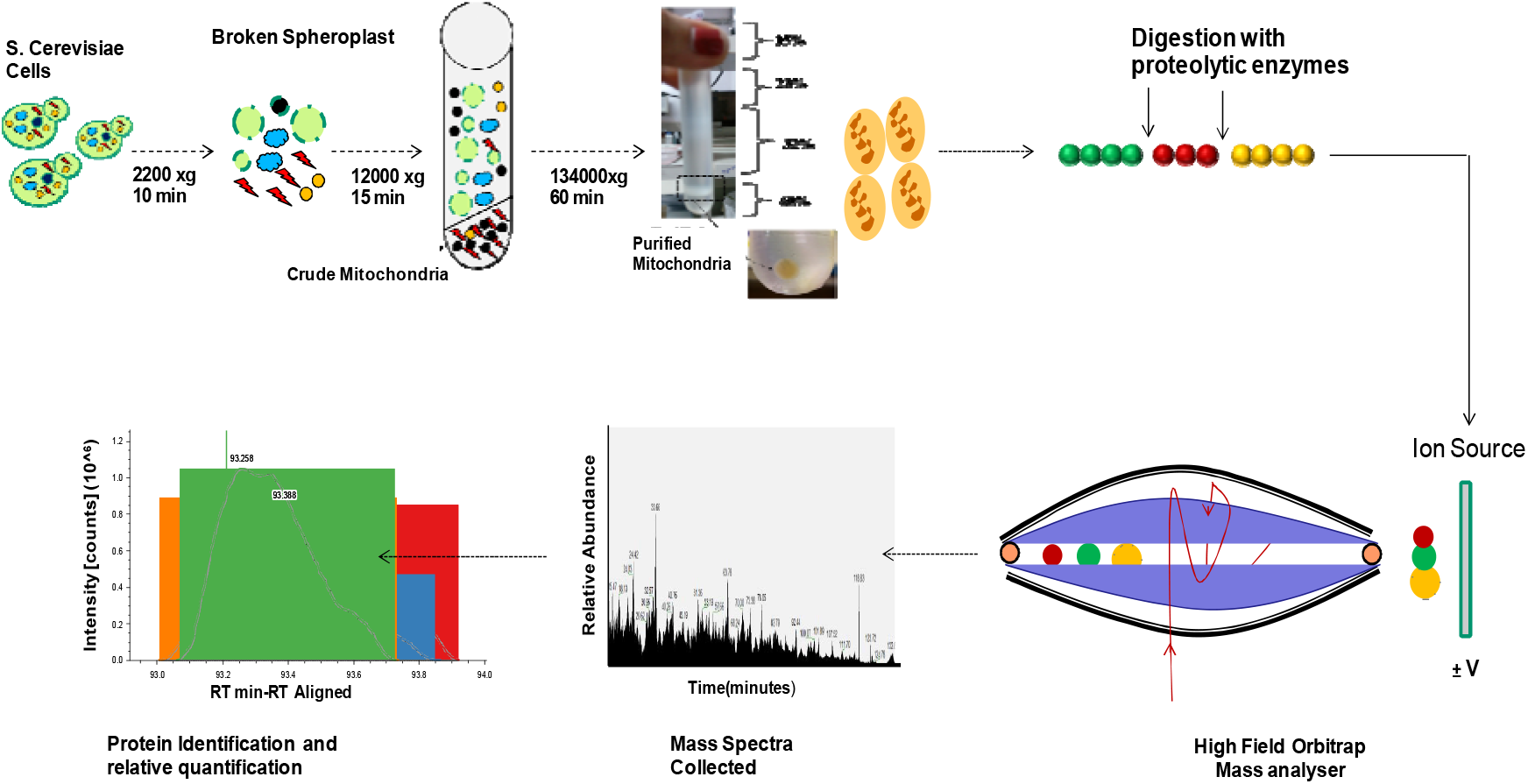
A schematic workflow applied for proteomic analysis. Mitochondria were isolated by sucrose density gradient centrifugation from *S.cerevisiae* (wtp53) cells. Mitochondrial proteins were isolated and digested with trypsin. Mitochondrial peptides were separated by mass spectrometry based on the mass-to-charge ratio (m/z) and further quantification was performed based upon combined peak intensity of MS spectru m.

#### G6PD enzyme activity

G6PD activity was determined as described previously (26). Cell lysates were added to the reaction buffer containing 50 mM Tris, 1 mM MgCl_2_, pH 8.1. First, the combined activity of G6PD and 6PGD was measured by the change in the rate of conversion of NADP^+^ to NADPH in the presence of substrate i.e. glucose-6-phosphate (G6P). In another cuvette, substrate 6 Phosphogluconate (6PG) was added to obtain only activity of the second enzyme. The rate of change was measured over 5 minutes of incubation period at 25°C. G6PD activity was calculated by subtraction of 6PGD activity from the total enzyme activity. For calculation of specific enzyme activity, rate of increase in absorbance at 341nm was divided by molar extinction coefficient of NADPH (6270). Enzyme activities were normalized based on protein concentration, which was determined by a Bradford protein assay.

#### Measurement of NADPH level

NADP^+^ and NADPH were measured based upon the absorbance of reduced coenzyme at 340nm by spectrophotometer (27). Samples were prepared in a defined buffer where three readings were taken, as the first reading represented the total amount of reduced nucleotides i.e. NADPH and NADH (A1). For the second reading, the cell extract is incubated with glucose-6-phosphate dehydrogenase (G6PD) to convert all the cellular NADP^+^ to NADPH (A2). The third reading was calculated based upon the decrease in the absorbance at 341nm after the total NADPH was converted to NADP^+^ by glutathione reductase (GR) i.e. (A3).A1-A3 represents the total amount of NADPH in the particular sample. The NADPH concentrations from unknown samples were determined by standard calibration curve that shows a linear increase in absorbance at 340nm with an increase in the concentration of NADPH from .001 to 0.1mM in one ml of the reaction mixture.

#### Mitochondrial colocalization and integrity

For staining mitochondria, we use Mitotracker^TM^ Red CMX Ros (Molecular Probes, Invitrogen) dye. WT and HXK2-GFP strains were grown at 30°C overnight in sucrose and glycerol lactate media, for p53 activation exponential phase cells were induced with galactose for 4-5 hours. Cells (0.1OD_600_) were stained with 100nm Mitotracker red (MTR-CMX Ros) 30°C for 45 min. The red fluorescence was visualised by confocal laser scanning microscopy and Flow cytometry.

#### Western Blotting

15μg of total cytosolic or mitochondrial protein was subjected to SDS PAGE. Then, the proteins were transferred from gel to nitrocellulose membrane. Immunodetection was performed with anti-p53/HA/G6PD/β-Actin/Porin antibody at dilution of 1:5000. The membrane was then subjected to alkaline phosphatase conjugated secondary antibody and protein bands were detected using solution that contained NBT, BCIP, 50mM MgCl_2_, 0.1M NaCl, 0.1M Tris-Cl buffer pH 9.5.

#### Measurement of intracellular ROS levels

Intracellular Reactive Oxygen Species (ROS) were detected by using the oxidant-sensitive probe 2’, 7’-dichlorodihydrofluorescein diacetate (DCDHF-DA). From the overnight grown culture of cells, OD_600nm_ of 0.1 was used further for induction with the galactose. After 6-8 hours of incubation, cells were harvested. H_2_O_2_ induction was used as positive control while water was added to the cells as a negative control. To an Equal number of cells, 20μm/l of DCDHF-DA was added in a shaking incubator at 28°C for 20 min. The cells were then washed and resuspended in 1 ml of 50 mM Tris-Cl buffer (pH 7.5). Two drops of chloroform and one drop of 0.1 % (w/v) SDS was added and the cells were vortexed for 20 sec. Incubated at room temperature for 15 min to allow the dye to diffuse into the buffer. Cells were pelleted and the fluorescence of the supernatant was measured using a spectrofluorometer with excitation at 490 nm and emission at 518 nm.

## Results

Mitochondria were purified from yeast cultures (wtp53) grown under four different conditions, including fermentable (sucrose) and non-fermentable (glycerol) with and without 2% galactose induction, as the former is required for over-expression of p53. All four mitochondrial samples were fractionated on SDS-PAGE followed by digestion with trypsin and analysed further by LC-MS/MS in triplicates (Two biological replicate and One technical replicate). To further confirm the complete recovery of peptides after In-gel digestion, In-Solution protein digestion was performed separately and proteomics results were compared where similar fold difference related to protein expression was observed. Quantification based upon peak intensities of MS signals across retention time with a defined mass window known as area under the curve (AUC) was used. Analyte intensity versus retention time profiles was generated from where the area under the curve or summed peak intensities is calculated. Relative peptide amount is proportional to the summed peak intensities.

### Coverage of Yeast mitochondrial proteome

A total of 1120 unique mitochondrial proteins with 1% FDR (false discovery rate) and more than 2 unique peptides were identified, indicating the involvement of mitochondria in various cellular processes (Supplementary Table-III). A volcano plot was generated to identify differential protein expression in tested conditions. It Plots fold-changes (x axis) and high statistical significance (−log10 of p value, y axis). When mitochondrial protein expression of sucrose grown cells after p53 over-expression was compared with its control (i.e. without p53 over-expression), from a total of 1120, 133 proteins were found to be differentially expressed. Out of 133, 69 proteins (upper right) were found to be up-regulated (P<0.05 and fold change of >1.5 fold), while 64 proteins (upper left) were found to be down-regulated (P<0.05 and a fold change of < 0.6 fold) in similar situation (Fig. 2a).

**Figure.2:**
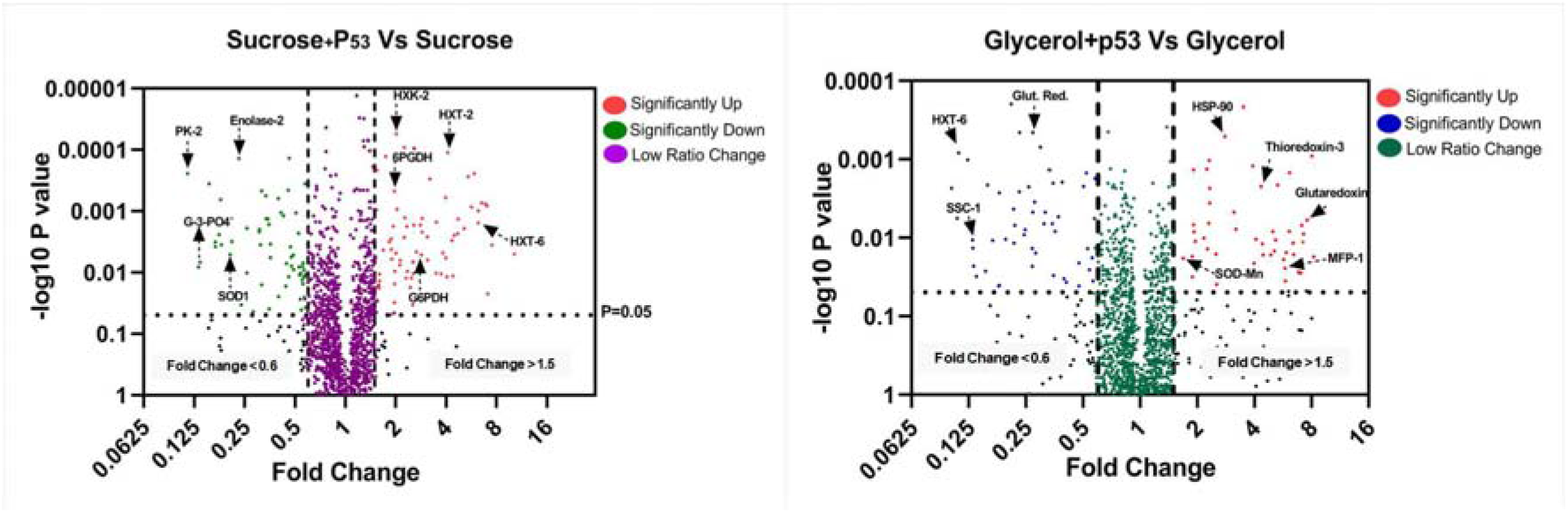
Volcano plot showing the statistical evaluation of the *S. cerevisiae* mitochondrial proteomic data: A volcano plot of the mitochondrial protein expression changes in (a) Sucrose grown cells after over-expression p53 expression (b) Glycerol grown cells after p53 over-expression when compared with their respective controls (without p53 over-expression). Data points in upper right denote significantly up (ratio >1.5 fold) and upper left denotes significantly down (ratio <0.6 fold) with P<0.05. The vertical axis indicates −log10 (p-value). The horizontal axis indicates log2 fold change, here labelled on the fold change scale. Probability values were derived by Student’s t-test. The volcano plot was generated using GraphPad Prism software version 6.01 for windows.

Similarly, when mitochondrial protein expression of p53 over-expressed glycerol-grown cells was compared with its control, out of 106 differentially expressed proteins, 56 were found to up-regulated (upper right) whereas 50 were found to be significantly down-regulated (upper left) (Fig. 2b). Summary of all differentially expressed proteins in both conditions is listed in the table (Supplementary Table-IV) and (Supplementary Table-V).

Of this list, a number of proteins that are up-regulated in p53 over-expressed conditions are belongs to a group that is involved in the antioxidant activity because oxidative stress is associated with p53-dependent cell cycle arrest; apoptosis and DNA repair (28). This data gave us the confidence that our proteomics analysis reflects the actual physiological change that is likely to occur in mitochondria.

Surprisingly, what we observed was that the members of the Pentose Phosphate Pathway (PPP) such as glucose-6-phosphate dehydrogenase, 6-phosphogluconate dehydrogenase, transaldolase, transketolase (Table-1) were found to be up-regulated in response to p53 in mitochondria isolated from sucrose grown cells. A similar increase was not observed in mitochondria isolated from glycerol-grown cells after over-expression of p53. Presence of PPP enzymes inside the mitochondria of *S.cerevisiae* was very intriguing and we decided to carry out a detailed analysis of this pathway in relation to p53 over-expression.

**Table 1:** Differential expression of Pentose Phosphate Pathway enzymes and hexokinases as identified by quantitative proteomics: Summary of mitochondrial proteins with increased expression of pentose phosphate pathway enzymes in p53 over-expressed sucrose grown cells when compared with its control and non-fermentable conditions, proteins were identified with >2 unique peptides and >1.5 fold Up-regulation (after 1%FDR). The values are mean ± SD of three independent experiments.

| Sr. No | Protein Name | UniProt<br>ID | Unique<br>Peptides | Coverage<br>(%) | Fold<br>Change<br>(B/D) | Sucrose<br>(A) | Sucrose<br>+gal (B) | Glycerol<br>(C) | Glycerol<br>+gal (D) | P value |
| --- | --- | --- | --- | --- | --- | --- | --- | --- | --- | --- |
| **Pentose Phosphate Pathway Enzymes** |  |  |  |  |  |  |  |  |  |  |
| 1 | Glucose-6-Phosphate Isomerase | P12709 | 11 | 12 | 1.36 | 81 | 132 | 88 | 97 | 0.0073 |
| 2 | 6- Phosphoglucolactonase | P53315 | 3 | 6 | 3 | 104 | 209 | 16 | 70 | 0.0089 |
| 3 | Glucose-6-Phosphate<br>Dehydrogenase | P11412 | 5 | 11 | 6.7 | 90 | 264 | 25 | 39 | 0.005 |
| 4 | 6-Phosphogluconate<br>Dehydrogenase | P53319 | 9 | 14 | 7.9 | 131 | 167 | 79 | 21 | 0.001 |
| 5 | Triose Phosphate<br>Isomerase | P00942 | 7 | 35 | 3.3 | 296 | 180 | 38 | 53 | 0.015 |
| 6 | Transaldolase | P15019 | 8 | 23 | 8.3 | 110 | 230 | 31 | 27.2 | 0.0092 |
| 7 | Transketolase-1 | P33315 | 10 | 14 | 5.4 | 153 | 193 | 17 | 36 | 0.0191 |
| 8 | Sedoheptulose-1,7-biphosphatase | P36136 | 3 | 12 | 1.5 | 168 | 78 | 96 | 55 | ns |
| **Hexose Transporter and Hexokinase** |  |  |  |  |  |  |  |  |  |  |
| 1 | HXT-6 | P39003 | 3 | 28 | 11.4 | 126 | 228 | 56 | 20 | 0.0051 |
| 2 | HXK-1 | P04806 | 11 | 34 | 2.7 | 256 | 212 | 50 | 77 | 0.0470 |
| 3 | HXK-2 | P04807 | 10 | 31 | 3.5 | 155 | 190 | 24 | 53 | 0.0001 |

Peptide count per protein or peptide peak area under the curve for mitochondrial G6PD was generated during chromatographic separation (Fig.3). Sample B i.e. Suc + Gal was having higher peak intensity and lower retention time (107.416) pointing towards higher content of mitochondrial G6PD. Sample D i.e. Gly+Gal was showing lower peak intensity with higher retention time (107.527) pointing towards highly reduced G6PD content.

**Figure. 3:**
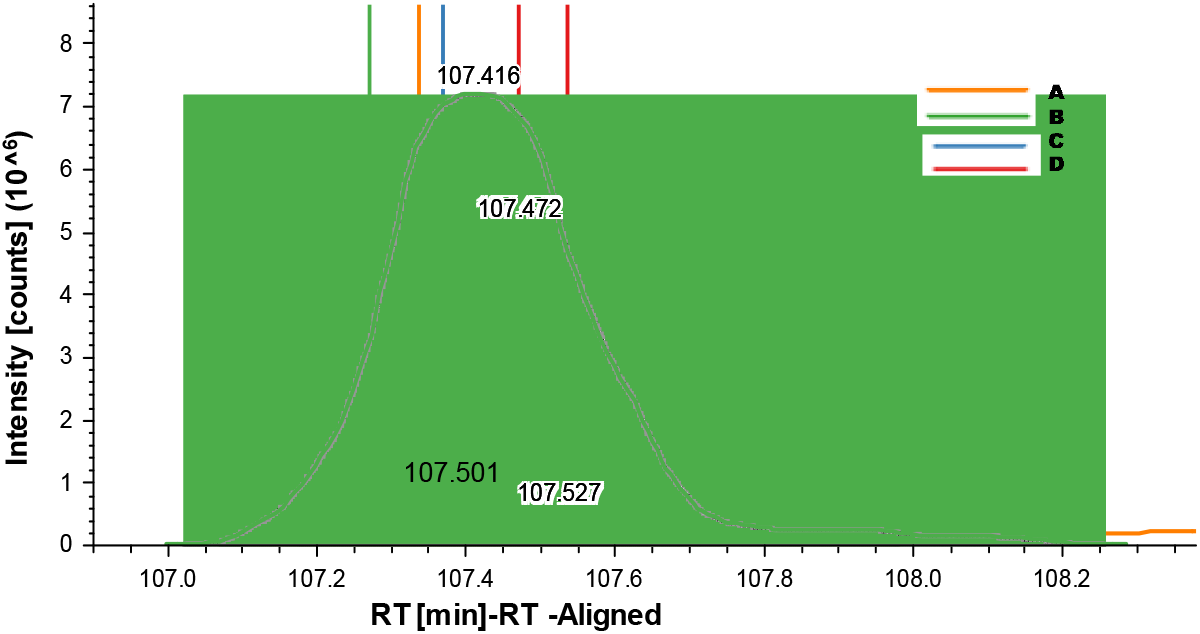
Quantification of mitochondrial G6PD based upon the comparison of the peak intensity of the same peptide in four tested conditions: Relative peptide amount of G6PD was plotted against retention time. Sample A, B, C and D are the mitochondria isolated from the cells grown in sucrose, sucrose+galactose, glycerol and glycerol+galactose respectively with a retention time of 107.416, 107.472 107.501 and 107.527. The quantification spectrum was generated using Proteome Discoverer (Thermo Fisher Scientific) and Mascot version 2.2.1 (Matrix Science) software.

### p53 induced apoptosis-resistant cells have increased expression of pentose phosphate pathway enzymes in their mitochondria

Based on the phenotypic observation that cells undergo apoptosis only when p53 is over-expressed in the cells grown in glycerol (non-fermentable) and not in the cells grown in sucrose (fermentable) (21), we hypothesized that the differential expression of mitochondrial PPP enzymes has a biological significance. Expression of p53 was confirmed in cytosolic as well as mitochondrial fractions isolated from cells grown in sucrose as well as glycerol after galactose induction (Fig.4).

**Figure 4:**
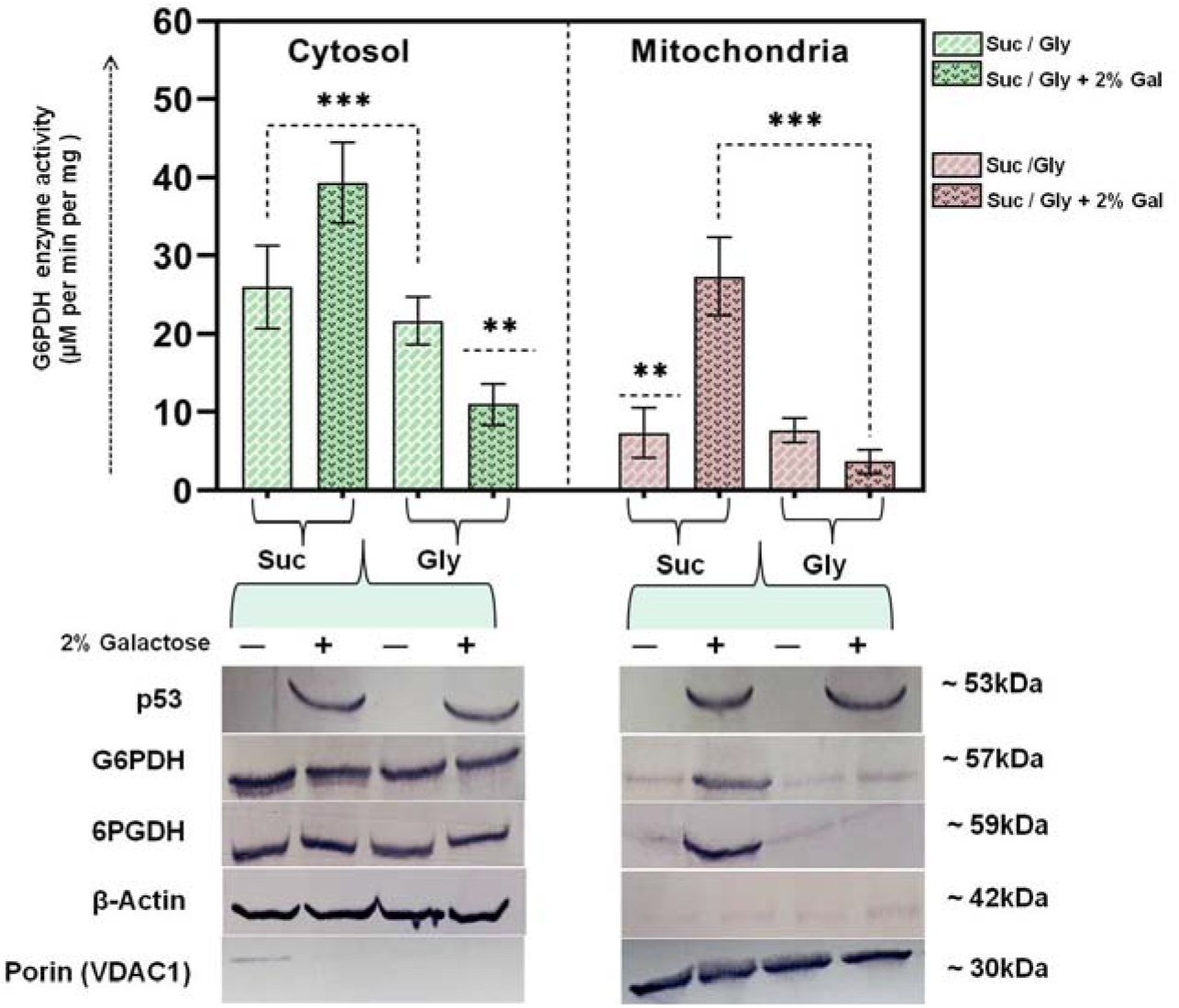
p53 regulates G6PDH activity and expression inside the mitochondria. G6PD activity (top) and protein expression (bottom) were analysed. G6PD activity (means ± s.d., n=3) in wt p53 cytosolic and mitochondrial samples treated with or without galactose. Cytosolic and enriched mitochondrial protein extracts were analysed by immunoblotting with anti p53, G6PDH and 6PGDH antibodies. Cytosolic marker β-actin (mABGEa) and mitochondrial marker VDAC were used as control.

To investigate the mechanism how p53 regulates PPP inside the mitochondria, we first monitored whether the increase in G6PD in mitochondrial proteomic analysis indeed reflects the physiological state of the mitochondria, we looked at the enzyme activity as well as the expression of G6PD by western blot analysis in both the cytosolic and mitochondrial fractions isolated from tested experimental conditions. Indeed both the cytosolic as well as mitochondrial G6PD enzyme activities were found to be increased up to ~2.5 fold upon p53 over-expression in the cells grown in sucrose in comparison to the cells without induction (Fig.4 above). Further, there was a noticeable reduction in cytosolic as well as mitochondrial enzyme activity in the cells grown in glycerol after p53 induction in comparison to their non-inducible counterpart.

To further confirm this increase in mitochondrial activity of G6PD, the level of protein expression was detected by western blot in both cytosolic as well as mitochondrial fractions. We were not able to detect any change in the protein level of cytosolic G6PD, surprisingly a variation in the mitochondrial content of G6PD was observed. Enhanced expression of mitochondrial G6PD was observed in the cells grown in sucrose after over-expression of p53 in comparison to its uninduced part. On the other hand, there was no significant difference in the expression of mitochondrial G6PD in glycerol and its inducible counterpart. To further confirm the role of p53 in regulating PPP, the second enzyme of PPP i.e.6 phosphogluconate dehydrogenase (6PGD) was targeted and checked for its expression under test conditions, where it followed a pattern similar to G6PD expression. Taken together, decreased expression of mitochondrial G6PD and other enzymes of PPP in response to over-expression of p53 in glycerol-grown cells enhance our understanding about its apoptosis-inducing activity.

Another important point to be noted is that a basal level of enzyme activity was also observed in mitochondria isolated from cells that are not induced with galactose probably suggesting that mitochondria inherently possess these enzyme activities, however less they are. While such observations have been reported both in yeast (29) as well as rat liver (30) but their physiological significance was not clearly revealed. Our observation that indeed increased PPP enzymes levels in mitochondria in response to p53 hints at a possible physiological role for this phenomenon.

If G6PD and 6PGD have any biological role, then the variable level of NADPH is expected in the mitochondria isolated from cells grown in sucrose under over-expression of p53 in comparison to the non-p53 expressed state. To determine this, NADPH levels were monitored in the cytosol as well as in mitochondria under all tested conditions. Strongly increased NADPH level was observed in mitochondria upon p53 over-expression in the cells grown in sucrose **~**up to 3 fold, (Fig.5a), support the ability of sucrose grown cells to escape from apoptosis even upon after p53 over-expression. The activity of G6PD was very low in the glycerol-grown cells over-expressed with p53, (Fig.4) and PPP might not contribute substantially to the overall NADPH production and hence most likely not able to reduce ROS. Deletion of G6PD further confirms the importance of PPP in preventing the sucrose grown cells from apoptosis as Δg6pd strain was showing a sharp decrease in NADPH content in both mitochondria as well as cytosol (Fig. 5b). This leads to abrogation in growth after p53 over-expression even in sucrose grown cells (Fig. 5c). Together these results show that increased levels of PPP enzymes and the resulting increased NADPH level inside the mitochondria prevent the sucrose grown cells from apoptosis even after p53 over-expression.

**Figure 5:**
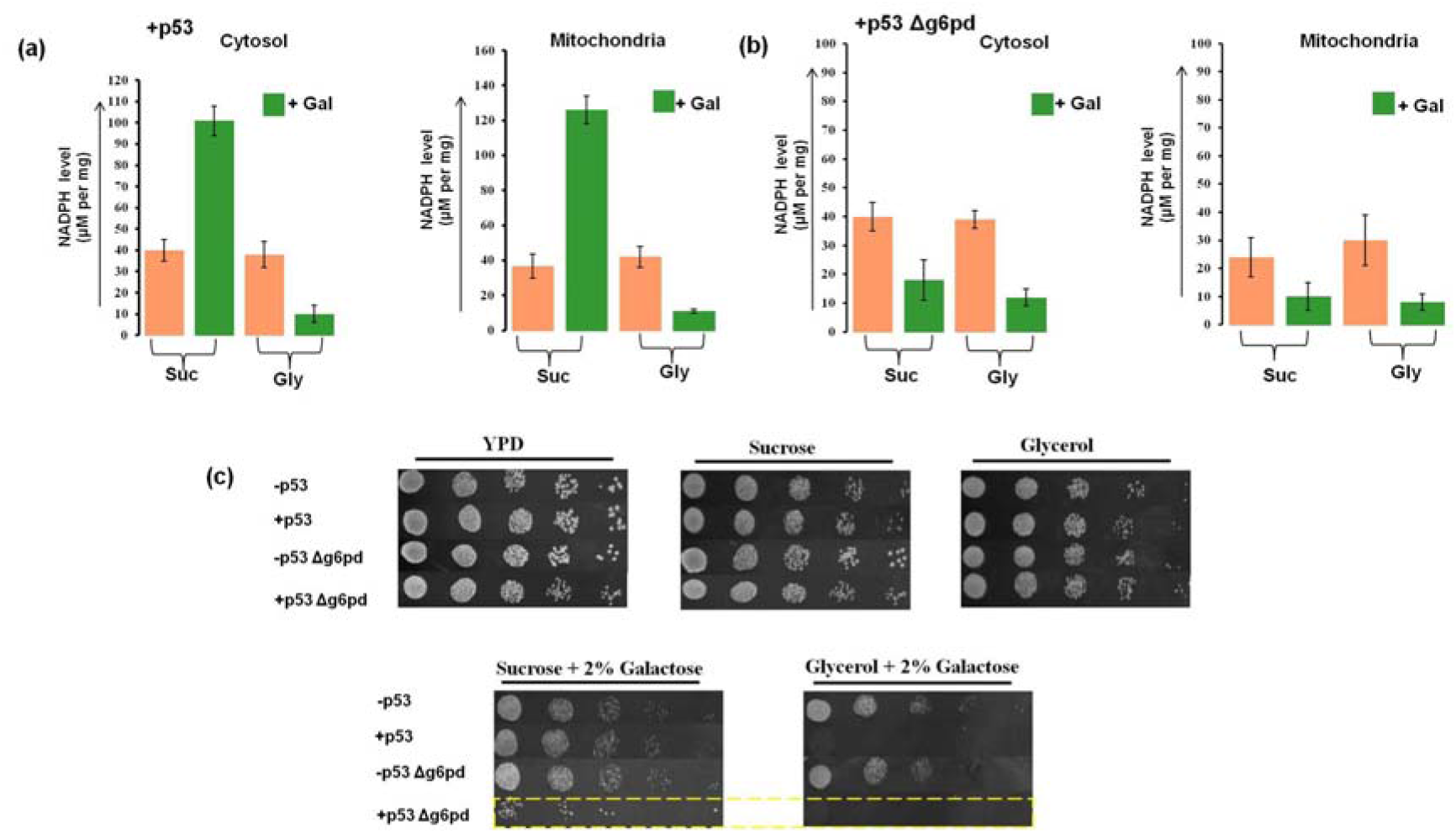
p53 regulates NADPH level inside the mitochondria through G6PD. (a), (b) NADPH level in +p53 and Δg6pd strain (means ± s.d., n=3) in cytosol and mitochondria separately in fermentable (sucrose) and non-fermentable (glycerol) with or without galactose induction. + Gal denotes induction with galactose in that particular sample. (c) The overnight grown culture of yeast strains without p53, +p53, and Δg6pd was diluted serially in sterile distilled water and 4μl of each sample was spotted on to medium containing sucrose or glycerol with (lower panel) and without 2% Galactose (upper panel). The growth phenotype was recorded after 48 hrs of growth at 30°C. Growth on Ypd was considered as a control.

### Over-Expression of hexose transporters as one of the determinants for Warburg effect

Over-expression of membrane high-affinity glucose transporters (Glut1and Glut3) is an important feature of several cancer cell lines, as their deletion has proved abrogated in-vitro tumour growth (31,32). No study related to the expression and significance of hexose transporters on mitochondrial membrane has been reported till now. Previous integrative analysis of mitochondrial proteome in yeast mentioned the presence of one of the hexose transporter i.e. HXT6 in their datasheet (33). However, the biological significance of this observation was not clear. Interestingly, we observed that the level of HXT6 was ~12 fold more in mitochondria isolated from p53 induced cells grown in sucrose in comparison to their p53 induced glycerol counterpart (Table-1, Supplementary Table-3). Fig. 6 shows an MS^2^ ion fragmentation spectrum for the selective peptide of HXT6. This spectrum shows a profusion of b and y ions depending upon N or C terminal fragmentation of the peptide “YLAEVGKIEEAK”. Reporter ion signal intensities belonging to the particular peptide are showing comparable difference pointing towards up and down-regulation of the protein in sucrose and glycerol respectively after p53 over-expression.

**Figure 6:**
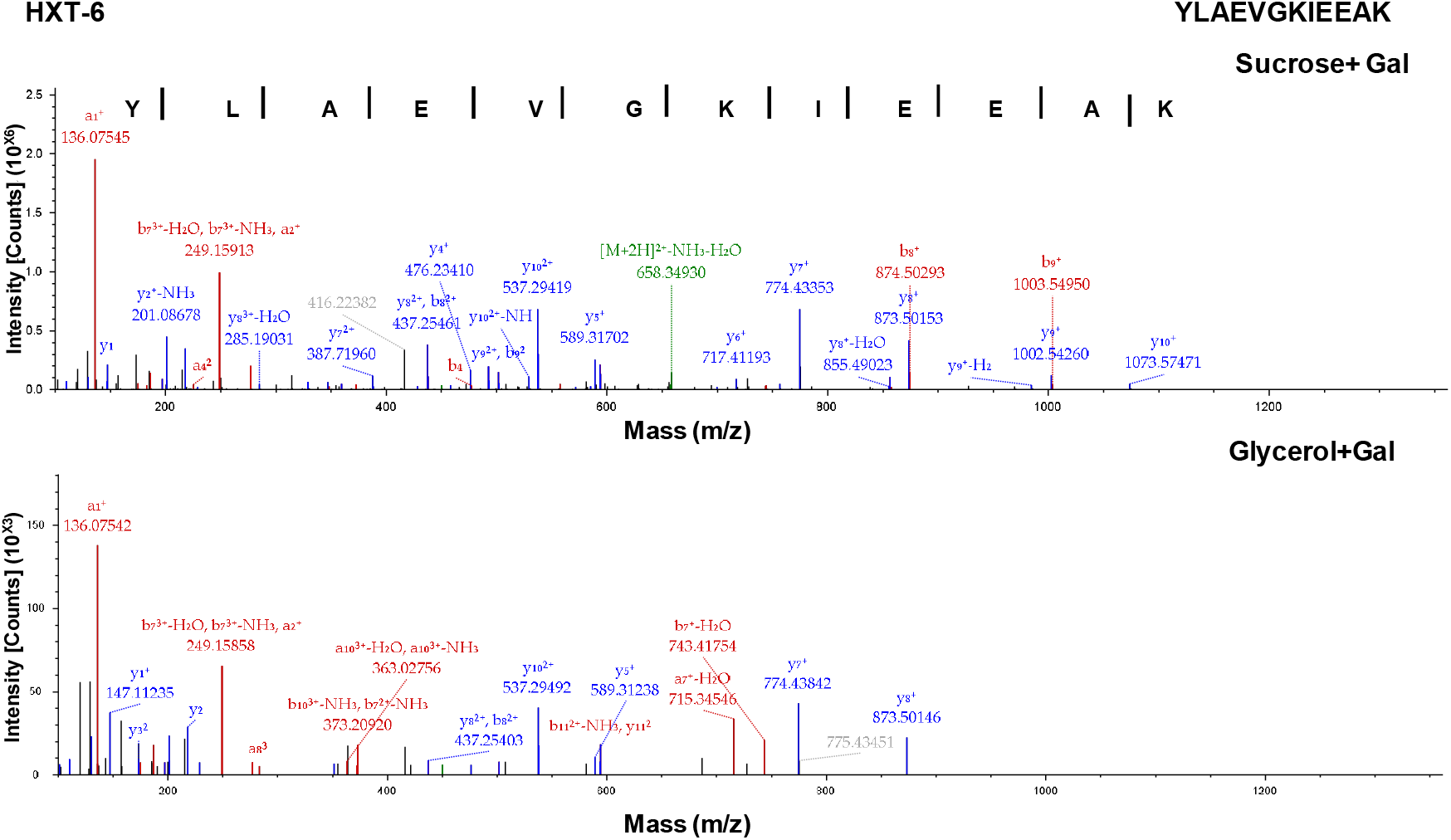
MS/MS spectra for identification and extraction of fragment ion chromatogram for peptide (YLAEVGKIEEAK) of HXT-6, showing differential expression in terms of intensity of particular ions. The signs b, y and a represent N-terminal type of fragments, C-terminal type of fragments or both type of fragments respectively. The fragmentation spectrum was generated using Proteome Discoverer (Thermo Fisher Scientific) and Mascot version 2.2.1 (Matrix Science) software.

If this increase is biologically relevant, we surmised that the deletion of HXT6 should not rescue the cell from p53 induced cell death. To validate this, HXT6 was deleted and the phenotype was monitored in different experimental conditions. Consistent with our hypothesis, deletion of HXT6 caused cell death in p53 induced cells grown in sucrose (Fig.7a). Taken together, the proteomics and drug susceptibility tests strongly pointed out that enhanced expression of HXT6 on mitochondria is critical for functional operation of mitochondrial PPP in sucrose grown cells after p53 over-expression.

**Figure 7:**
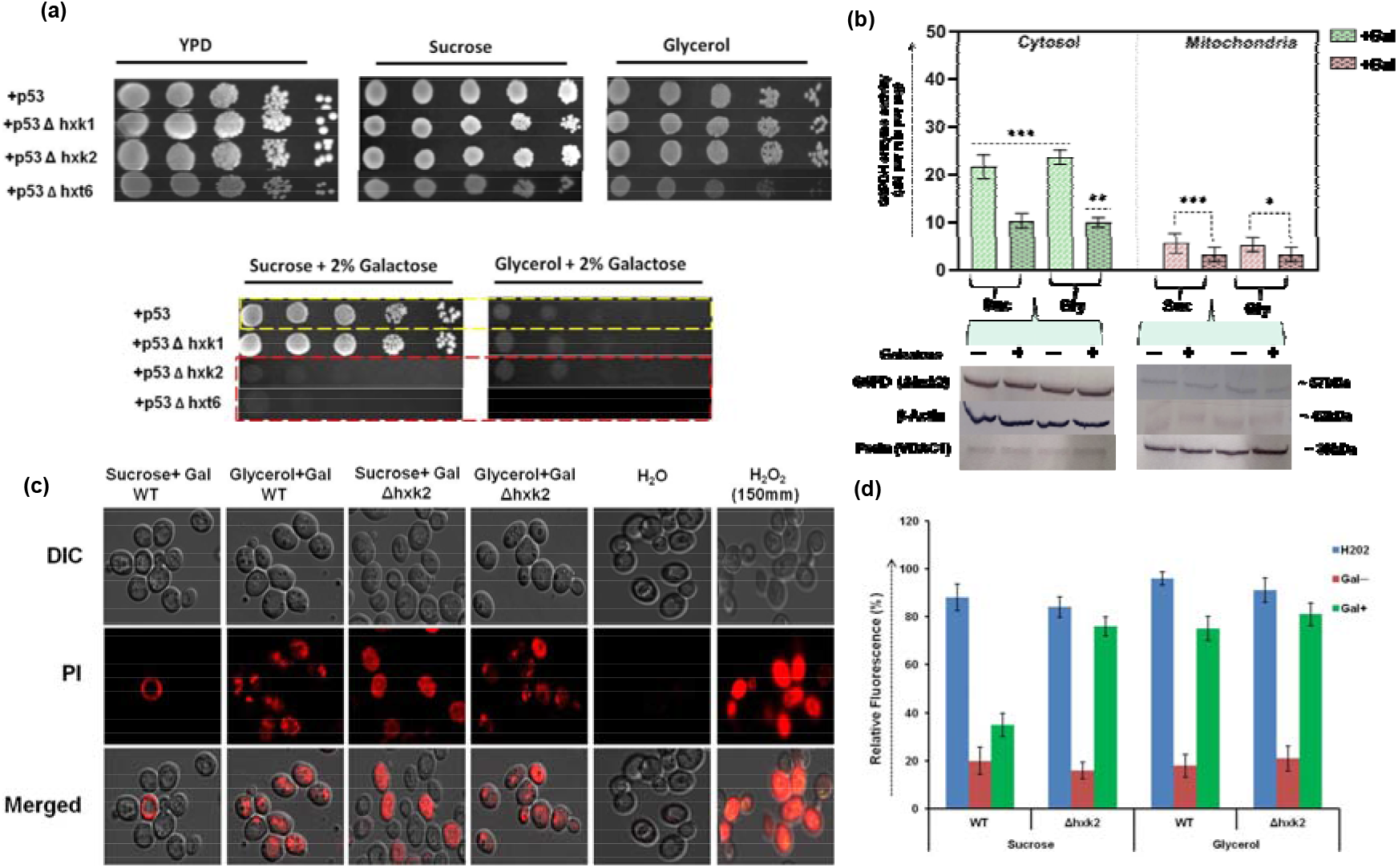
Deletion of HXK-2 exhibits impaired localisation of G6PDH to the mitochondria. (a) The overnight grown culture of yeast strains expressing p53, Δhxk-1, Δhxk-2, and Δhxt-6 was diluted serially in sterile distilled water and 4μl of each sample was spotted onto medium containing sucrose or glycerol with (lower panel) and without 2% Galactose (upper panel). The growth phenotype was recorded after 48 hrs of growth at 30°C. (b) G6PD activity (top) and protein expression (bottom) were analysed. G6PD activity (means ± s.d., n=3) in Δhxk2 cytosolic and mitochondrial samples treated with or without galactose. Cytosolic and enriched mitochondrial protein extracts were analysed by immunoblotting, cytosolic marker β-actin (mABGEa) and mitochondrial marker VDAC were used as control. (c) WT and Δhxk-2 cells were stained with 5μg ml^−1^ PI for 20 mins in dark at room temp. PI fluorescence was examined under the fluorescence microscope at λex = 480 nm from at least 300 cells in each experiment. (d) ROS generation was determined using DCDHF-DA (20 μmol/l, 15 min, 37°C).

### Deletion of HXK2 attenuates the influence of G6PD on p53 induced apoptosis

It has been known for long that HXK2 isoform is associated with mitochondrial fraction and has a role in preventing cancer cells from entering into apoptosis (20). Therefore, we wanted to know whether HXK2 plays any role in preventing apoptosis induced by over-expression of p53 in yeast. Our mitochondrial proteomic analysis also reflected the differential pattern of HXK2 expression in all four tested conditions.

For this, we first deleted HXK2 and observed its growth in the presence of sucrose or glycerol with and without p53 over-expression. As a control, we deleted HXK1 and subjected this strain to similar experimental analysis. It is clear from the growth phenotype, that the loss of HXK2, not HXK1 confers defective growth upon p53 over-expression even in the sucrose grown cells, suggesting that HXK2 is required for normal tolerance to induced p53 in sucrose grown cells. Consistent with this, reduced G6PD activity and no detectable presence of G6PD observed in the mitochondrial fraction isolated from p53 over-expressed sucrose grown cells of Δhxk2 cells as confirmed by western blot (Fig.7b).

Since ROS plays an important role in apoptotic cell death, fluorescence-based measurement of mitochondrial ROS was performed using an oxidant-sensitive probe (DCDHF-DA) in Δhxk2 cells along with (wtp53) strain under all previously tested conditions. After p53 over-expression in sucrose grown cells of Δhxk2 strain, ~3fold increase in ROS accumulation was observed inside the mitochondria in comparison to wild type under the same condition (Fig.7c). Cell death induced by p53 in the cells grown in sucrose after deletion of HXK2 was confirmed by Propidium Iodide (PI) staining, as PI-positive cells were confirmed dead (Fig.7d). Cells treated with 150mm H_2_O_2_ were used as a positive control whereas cells treated with water were used as a negative control.

### HXK-2 GFP is normally a cytosolic protein but colocalizes with mitochondria after over-expression of p53 in Sucrose grown cells

We constructed a C-terminus GFP tagged HXK2 protein, expressed from plasmid pYM28 and monitored translocation of this protein in the cells grown in sucrose and glycerol, with and without p53 expression to determine whether, the protein enters to the mitochondria or not. The functionality of HXK2-GFP strain was confirmed by spot dilution assay in sucrose and glycerol media with and without galactose, where it behaves similar to WT strain (data not shown). For mitochondrial staining, we used a mitochondria-specific vital dye, Mito Tracker™ Red that is known to passively diffuse across the yeast cell membrane and accumulate in active mitochondria that can be visualised by enhanced fluorescence after its oxidation in respiring mitochondria.

We observed that in the cells grown in sucrose, green fluorescent protein-tagged HXK2 was found diffused throughout the cytosol and partially colocalises with mitochondria (Fig.8a). After over-expression of p53 in the sucrose grown cells, we observed a clear detection of protein attached to the mitochondria as determined by co-staining with Mito Tracker^TM^ CXMRos (Fig.8a).

**Figure 8:**
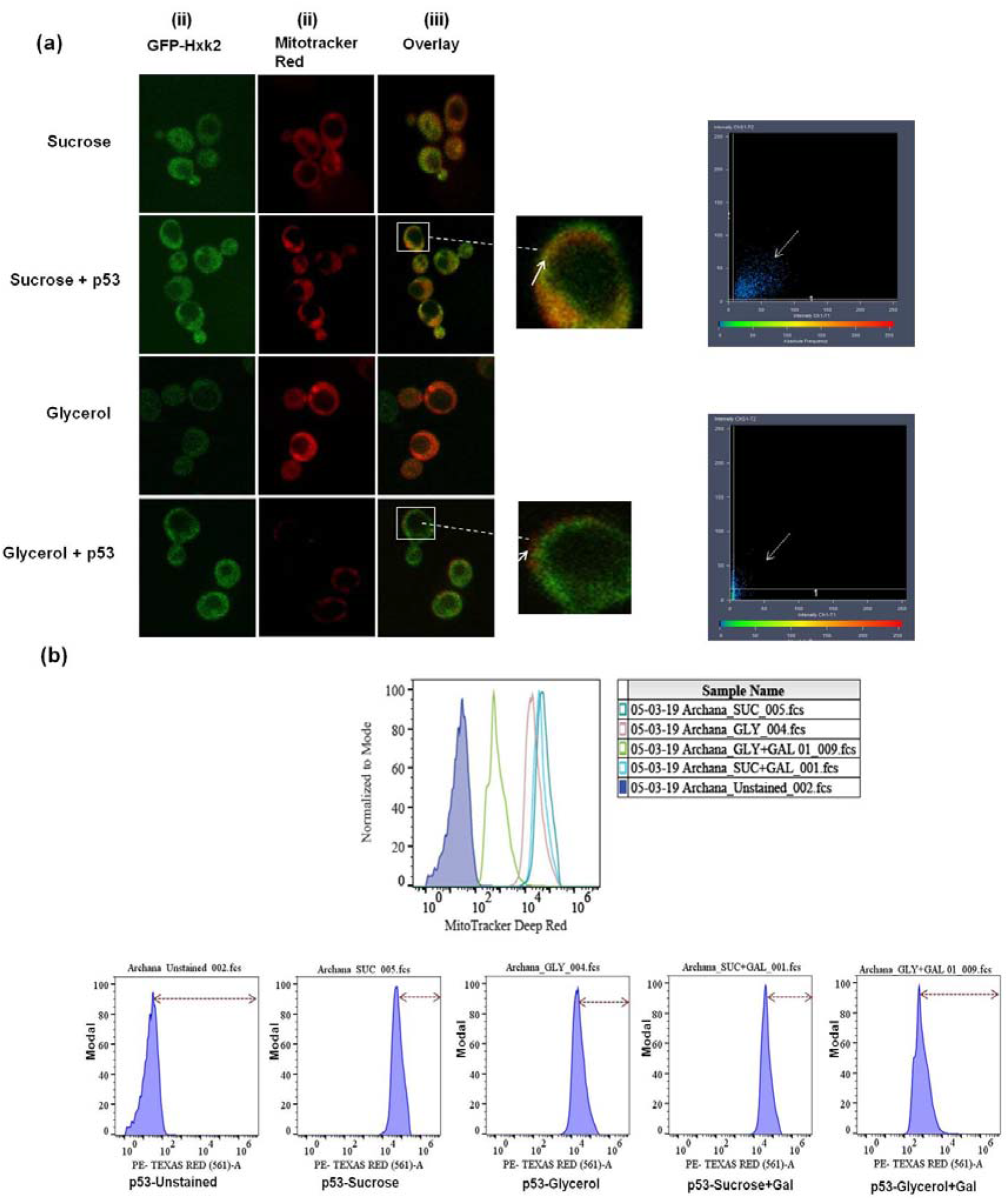
Hxk2-GFP Colocalize with mitochondria after p53 over-expression in Sucrose grown cells. (A) *S.cerevisiae* (+p53) cells transiently expressing GFP-HXK2 were treated with Mitotracker Red CMXRos (100nm) to stain for mitochondria and then examined after 40min by laser fluorescence confocal microscopy. The field shown was independently visualized at the appropriate wavelength for GFP (λex=633nm), Mitotracker Red CMXRos (λex=405nm) and the two images were then overlaid. Bar 10μm. (B). Representative flow cytometry histogram with an overlay of all 4 tested conditions along with unstained as control showing the difference in Mitotracker red expression

Reduced HXK2 expression was observed in the cells grown in glycerol (Fig.8a), as observed previously also (34). In spite of reduced HXK2 expression, mitochondria are fully functional which can be visualized by enhanced fluorescence of Mito-tracker red. However, the poor red fluorescence intensity was observed after over-expression of p53 in glycerol-grown cells pointing towards compromised mitochondrial functionality as the uptake of Mito Tracker CXMRos is mitochondrial membrane potential-dependent (35). It was further reaffirmed when we examined the mitochondrial integrity with FACS analysis by employing mitotracker red dye (Fig.8b). Cells grown in Glycerol+galactose are having histogram shift towards unstained pointing towards reduced staining with mitotracker red in comparison to other samples. These results indicate that co-localisation of HXK2 to the mitochondria is necessary for the operation of functional PPP inside the mitochondria and the resulting increase of NADPH.

## Discussion

One of the fundamental issues is how cancer cells manage to escape from the onslaught of superoxide radicals that induce apoptosis. Although the results presented here are obtained using *Saccharomyces cerevisiae*, we believe that in all probability a similar mechanism could be operative in mammalian cells. This confidence is based on a large number of studies conducted in yeast that has many parallels in humans (29, 36, 37, 38, and 39). G6PD catalyzes the first step of the pentose phosphate pathway (or hexose monophosphate shunt) to provide ribose for nucleotide metabolism and NADPH as a reductive substrate (40). In this study, we identified an enhanced expression of G6PD in the mitochondria of sucrose grown *S.cerevisiae* cells after over-expression of p53. Western blot from pure mitochondrial fraction, G6PD activity assay and proteome analysis clearly pointed towards mitochondrial localisation of G6PD. Campbell and Berenofsky have made a similar observation in 1978 when they partially purify the G6PD from yeast mitochondria and found properties similar to its cytosolic isoform in terms of electrophoretic mobility, isoelectric point, molecular size and Km for NADP^+^ and glucose-6-P (29).

Our experiments demonstrate that level of HXK2 bound to mitochondria greatly exceeds in p53 over-expressed sucrose grown cells in comparison to non-expressed counterpart. This step seems to be crucial in the translocation of the PPP enzymes into mitochondria after p53 over-expression. So far, only one report has demonstrated the presence of PPP enzymes in the rat liver mitochondria (41). It is reported that when glucose flux is high, the mitochondria depends on NADPH produced by G6PD (42). The present study indicates that increased mitochondrial G6PD is a significant provider of NADPH, an important component of mitochondrial antioxidant defence system against ROS because mitochondria under p53 over-expressed conditions are affected with higher ROS and reliant on the pool of NADPH to maintain the reductive environment inside the mitochondria. Our study for the first time brings out the possibility that escape from apoptosis is triggered by mitochondrial glucose-6-P flux mediated through HXK2, as its deletion confirmed reduced NADPH level (Fig.7), increased mitochondrial ROS (Fig.7) and apoptosis (Fig.7).

Interestingly, cells went for apoptosis after over-expression of p53 in non-fermentable carbon source. Our results demonstrate that in glycerol-grown cells only basal level of HXK2 expression was observed, which is not sufficient to attach and flux glucose-6-P towards ROS affected mitochondria that ultimately leads to cell death. Hence, carbon source-dependent expression of HXK2 regulates the operation of functional PPP inside the mitochondria only on p53 over-expression. Enhanced expression of HXT6 transporter on the mitochondrial membrane of sucrose grown cells (Table-1, Fig.3), pointing towards its importance in relation to the supply of glucose for functional operation of PPP.

If HXK2 is present on the outer membrane and does not enter mitochondria, how Glucose 6-phosphate is made available as a substrate for G6PD is not clear. It can be possible that the Glucose-6-P generated outside the mitochondria is transported into the mitochondria through VDAC. This is based on the reports that VDAC and HXK2 are in close association in both *S. cerevisiae* as well as humans (20,43). Alternatively, HXK2 could get into mitochondria to provide Glucose 6 Phosphate to G6PD. If this idea is true, then the presence of HXT6 makes sense. If the former possibility is true, the role for HXT6 on the mitochondrial membrane is not clear. In our results, we consistently observe the presence of the minuscule amount of G6PD in mitochondria samples isolated from cells not over-expressed for p53. Thus, it is possible that may be mitochondria have this enzyme inherently (Fig.4), but its translocation gets accentuated only when cells are exposed to ROS. Presence of full complement of pentose phosphate pathway enzymes inside the mitochondria may function as a source of ribose-5-P for the synthesis of mitochondrial nucleotides. No significant change in the expression level of Isocitrate dehydrogenase (ICDH), another NADPH forming mitochondrial enzyme was observed between p53 expressed and not expressed conditions in both fermentable and non-fermentable conditions (Table-2).

**Table-2:**
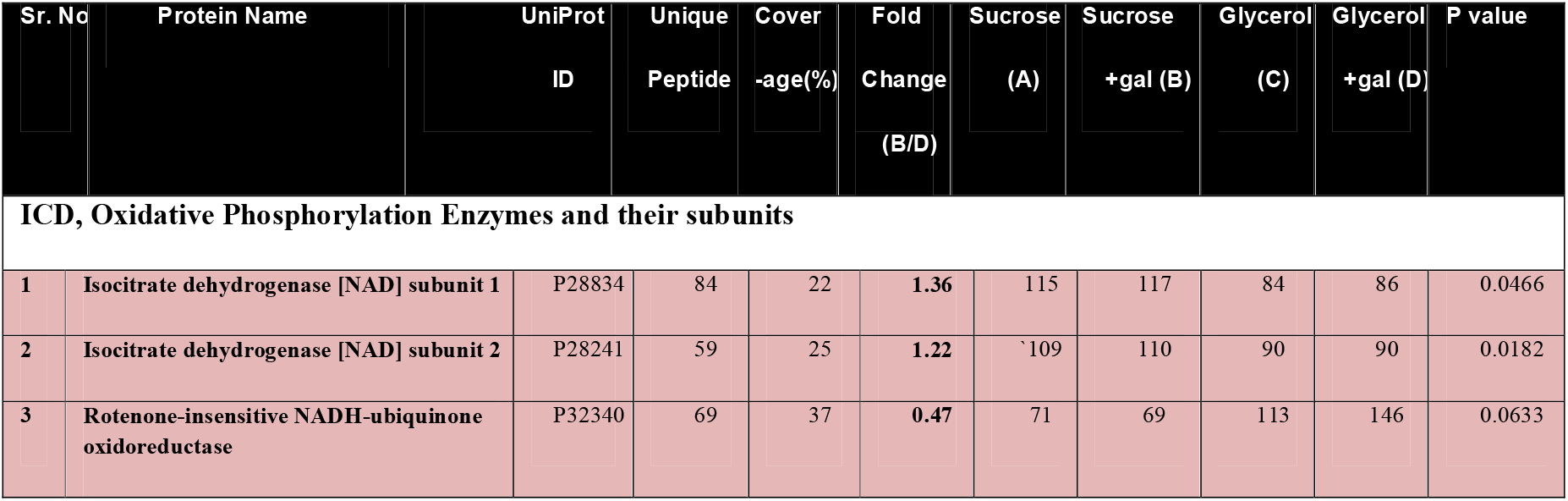
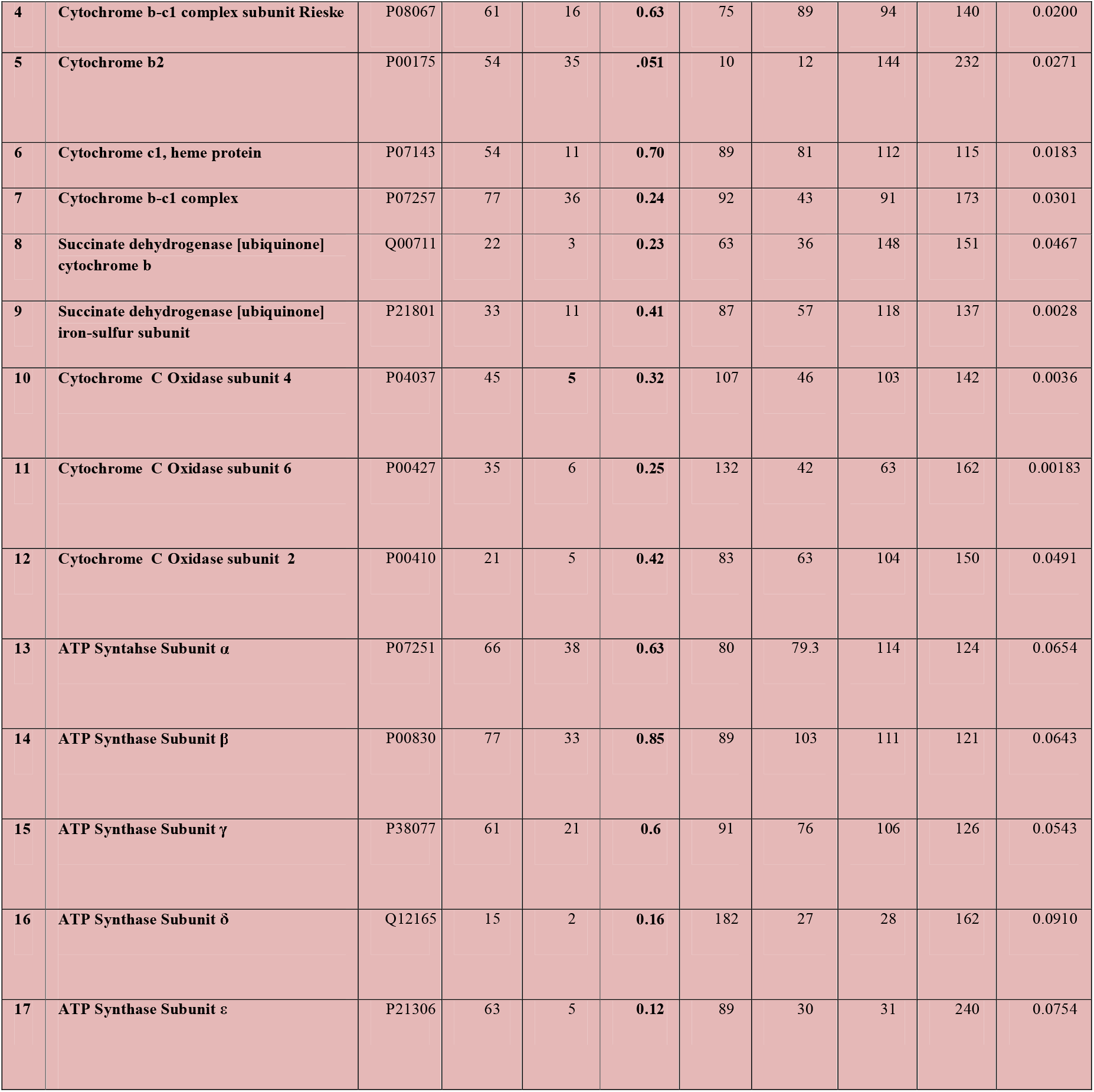
Differential expression of Isocitrate Dehydrogenase (ICDH) and Oxidative Phosphorylation enzymes as identified by quantitative proteomics: Summary of mitochondrial proteins with decreased expression of oxidative phosphorylation enzymes in p53 induced sucrose grown cells when Compared with control and with non-fermentable conditions. 17 down-regulated proteins expressed less than <0.6 fold (<1%FDR). The values are mean ± SD of three independent experiments.

Enhanced glycolytic activity and impaired oxidative phosphorylation (Warburg effect) is a widely accepted phenotype of tumour cells (1). So we decided to check the expression level of enzymes (and their related subunits) involved in oxidative phosphorylation inside the mitochondria of yeast cells where PPP is upregulated after over-expression of p53. Interestingly, reduced expression of enzymes (NADH ubiquinone oxidoreductase, Succinate Dehydrogenase, Cytochrome b-c1 complex, Cytochrome C oxidase, ATP synthase) was observed in p53 over-expressed sucrose grown cells in comparison to control.

In conclusion, the human p53 seems to be functional in *S.cerevisiae*; it could translocate to mitochondria, generate reactive oxygen species, disturb mitochondrial membrane potential and ultimately cause cell death. In our study, we observed a novel escape mechanism of yeast from apoptosis after over-expression of p53. Operation of functional mitochondrial PPP pathway reduced oxidative phosphorylation, an enhanced association of HXK2, HXT6 to the mitochondrial membrane, increased mitochondrial NADPH content restores yeast growth and escapes it from apoptosis. From this metabolic point of view, we can say that fermenting yeast and tumour cells share several features related to Warburg and Crabtree effect. This metabolic control is a key element of apoptotic escape and tumour progression. An impressive approach would be to study this fermenting yeast apoptotic evasion strategy in mamalian cancer cell lines.

## Supporting information

III

## Acknowledgement

Archana Kumari Redhu is thankful to the Department of Bio-Science and Bio-Engineering, Indian Institute of Technology-Bombay for awarding Post Doctoral Research Fellowship. For Proteomics facility authors would like to thank MASSFIITB facility supported by Department of Biotechnology (BT/PR13114/INF/22/206/2015). A.K.R. would like to thank Mr Vipin Kumar (Proteomics Facility) for his valuable and fruitful discussions regarding proteomics data.

